# Flexible hippocampal representation of abstract boundaries supports memory-guided choice

**DOI:** 10.1101/2024.07.23.604745

**Authors:** Mariachiara Esposito, Lubna Abdul, Ameer Ghouse, Marta Rodriguez Aramendía, Raphael Kaplan

## Abstract

Hippocampal cognitive maps encode the relative locations of spatial cues in an environment and adapt their representation when boundaries geometrically change. Hippocampal cognitive maps can represent abstract knowledge, yet it’s unclear whether the hippocampus is sensitive to changes to the extreme coordinates, boundaries, of abstract spaces. We created a memory-guided choice task to test whether the human hippocampus and medial prefrontal cortex(mPFC) flexibly learn abstract boundary representations in distinct two-dimensional(2D) knowledge spaces. Participants built up a 2D map-like representation of abstract boundaries, where the hippocampus and mPFC represented a decision cue’s Euclidean distance to the closest boundary. Notably, mPFC distance representations selectively reflected individual performance improvements during the task. Testing for neural sensitivity to boundary-defined contextual changes, only the hippocampus flexibly represented abstract boundaries, which related to choice behavior. These findings suggest that abstract knowledge retrieval within dynamically changing contexts is facilitated by generalized mPFC and flexible hippocampal boundary representations.

## Introduction

The limits of our environment help structure how we think. Physical boundaries, independent of whether they are directly perceived [1] or imagined [2], are known to help guide spatial learning. Boundaries serve as a fundamental spatial orientation cue that guides learning in a host of species including humans [3–6]. In parallel, the discovery of spatially-modulated cells in the hippocampal formation [7] that are sensitive to the shape, distance, and direction to environmental borders [8] has bolstered the idea that hippocampal neurons form a cognitive map of the spatial layout of an environment [9]. Notably, a key feature of hippocampal cognitive maps is their ability to dynamically adapt to geometrical changes in environmental boundaries [10–17]. This dynamic adaptation is thought to facilitate flexible contextual processing in the service of spatial memory formation [18–23]. Taken together with theories that hippocampal cognitive maps extend to episodic memory storage [9, 24], these ideas implicate cognitive maps as an ideal format for flexibly representing abstract knowledge [25–27].

Building on these popular theories, mounting evidence in a variety of species has found that abstract knowledge is assimilated via cognitive map-like representations in the hippocampal formation [28–35, see 27, 36 for reviews] and other regions, namely medial prefrontal/orbitofrontal cortex(mPFC/OFC) [26, 37, 38]. More specifically, human hippocampus and mPFC fMRI signals reflect the Euclidean distance between learned stimuli during decision making [31], providing support for the importance of hippocampal-prefrontal map-like representation of abstract knowledge [26]. Yet, whether hippocampus or mPFC representations are sensitive to geometric changes to the ‘boundaries’ or extreme coordinates of abstract knowledge spaces is unclear. Still, the ability of cognitive maps to flexibly code abstract knowledge raises the possibility that abstract boundaries–continuous(e.g., price range) or categorical(e.g., types of cars) limits–could help guide map-like learning of abstract knowledge. Here, we test the hypothesis that map-like representations of knowledge in the human hippocampus and mPFC are sensitive to abstract boundary changes. In particular, we were interested whether 2D Euclidean distances to abstract boundaries were represented in the hippocampal-prefrontal circuit and if different abstract boundary-defined decision contexts could be distinguished by hippocampal fMRI signals. Addressing whether abstract boundaries are processed by the human hippocampus and mPFC during memory-guided choice in a similar way as physical borders during learning, we developed a two alternative forced-choice(2AFC) fMRI task. In the task, participants made similarity judgments on either the price or freshness of various fruit and vegetable goods without receiving any feedback.

Unbeknownst to participants, the continuous price and freshness variables for each produce good formed two-dimensional(2D) abstract spaces, with four of the goods lying at the extreme coordinates(boundary goods), one good fixed near the center(landmark good), and sixteen cued goods that were located within the boundaries. Inspired by human spatial memory experiments that have participants learn the location of stimuli and then cue them to remember where that stimuli is located in an environment, the fMRI task consisted of blocks containing encoding and decision components in the abstract space (Fig. 1A). Differing from the typical spatial memory experiment that has participants first remember cued goods and relate their location to visible boundaries and landmarks, we had participants remember boundary and landmark abstract good coordinates and then relate those values to a visible cued good.

**Figure 1:**
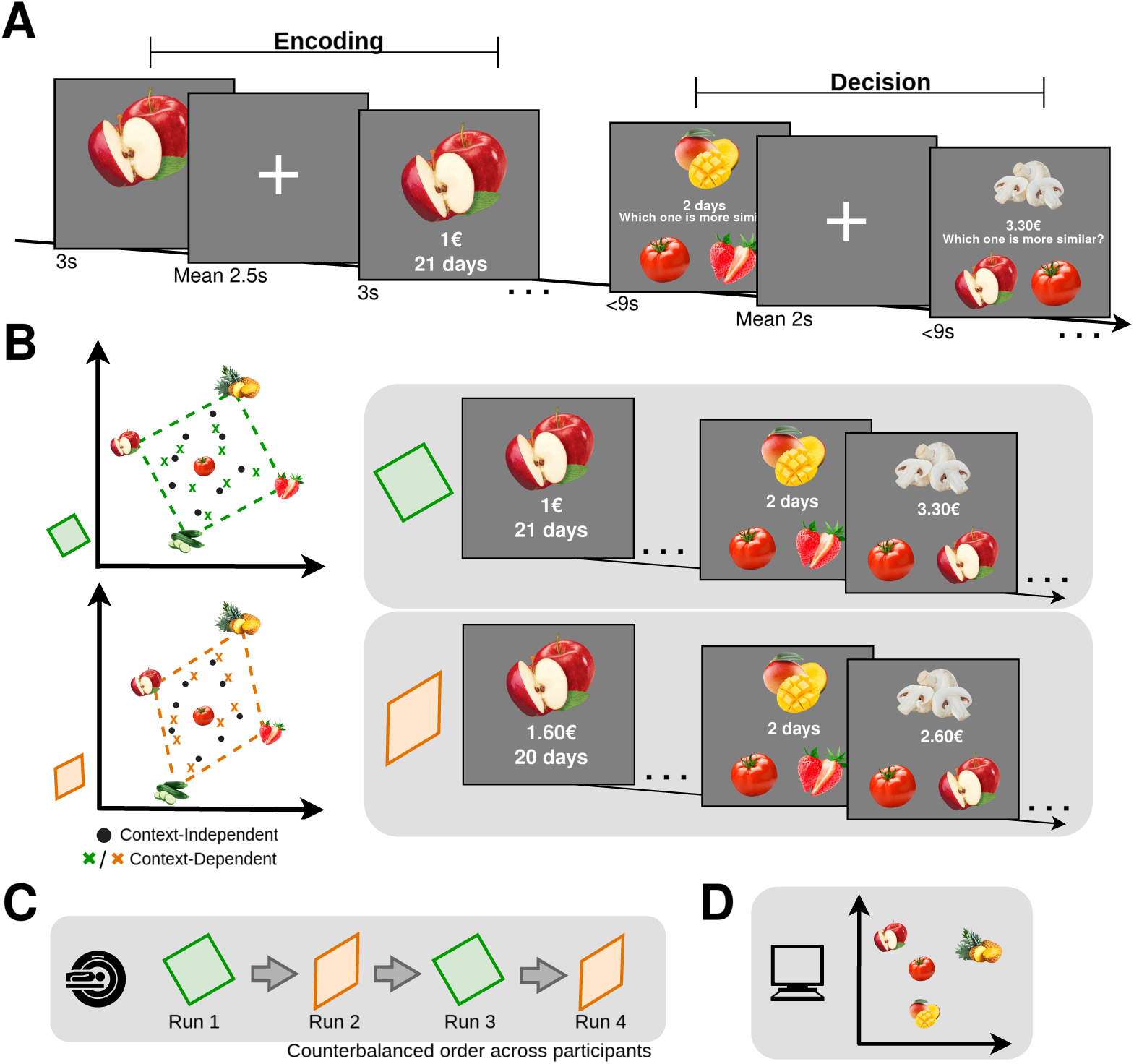
Experimental paradigm. **A**. fMRI Task. During encoding, participants were presented with continuous price and freshness variables for each of the boundary and landmark goods. Landmark and boundary goods were first presented without visible decision variables for 3s and then with visible decision variables for another 3s following a jittered intertrial interval(ITI mean=2.5s). Decision trials featured one of sixteen cued produce goods inside the space, and participants made a self-paced two alternative forced-choice(2AFC) based on whether that cued good’s price or freshness was more similar to that of the landmark, or its most proximal boundary good. No feedback on choices was provided during the fMRI task. The cued good’s relevant decision variable was visible on the screen throughout the trial, while the landmark and boundary variables weren’t shown during decision trials. **B**. Unbeknownst to participants, price and freshness variable coordinates for the four boundary goods formed two abstract shapes (the “square” shape and “distorted” shape in green and orange respectively), where the boundary goods are located at the extremes of the space and have two different sets of coordinates for each space. In contrast, the landmark good kept its position close to the center. The sixteen cued goods were located within the boundaries. Half of the cued goods kept the same coordinates in both square and distorted shape (context-independent goods), while the other half changed their positions based on the shape they were in (context-dependent goods). In an fMRI block, participants were first prompted to encode boundary and landmark good decision variables and then made similarity judgments relative to a cued good immediately following encoding. **C**. Participants performed each shape twice, alternating between square and distorted shapes, completing a total of four runs. The presentation order was counterbalanced across participants. **D**. After the fMRI task, participants performed a surprise drag-and-rate task outside the scanner. Participants were instructed to During encoding, the boundary goods and the landmark good were first presented without visible decision variables, and after an intertrial interval(ITI), were individually cued with their respective price and freshness variables at the same time. Immediately following their encoding of the landmark and boundary goods, participants were presented with the price or freshness of a cued good and chose whether it was more similar along that dimension to either the landmark good or the most proximal boundary good. The most proximal boundary good in each trial was selected based on its 2D Euclidean distance to the presented cued good. During 2AFC trials, the cued good was the only stimulus with a visible price or freshness variable. Crucially, the boundary produce goods had two distinct sets of price and freshness variable coordinates depending on which run they were featured. Given the increased cognitive demand of changing invisible abstract contexts during the middle of an fMRI run compared to spatial navigation tasks, each run was dedicated to one of two distinct boundary-defined abstract spaces. In contrast, the landmark coordinates were consistent in the two spaces(Fig. 1B). Each fMRI run included four blocks. Participants completed a total of four runs in the scanner, alternating between square and distorted shaped contexts(Fig. 1C).

Presenting participants with 2D coordinates for boundary/landmark goods to remember and then prompting them to make decisions on one variable/dimension allowed us to investigate two types of abstract distance: the distance that participants directly judged during the choices(e.g., the relative choice distance based on 1D price/freshness), and the Euclidean distances based on the variables they learned during the encoding phase. We cued participants with 2D variables to avoid overtraining, but participants crucially didn’t need both coordinates to accurately calculate the 1D relative choice distance. This design aspect allowed us to confirm with a general linear model(GLM) if the 1D distance and 2D Euclidean distance of a choice were both significant predictors of choice accuracy and test whether this predictability changed over the course of the experiment. If map-like learning of abstract knowledge spaces related to behavioral performance, the predictability of the Euclidean distance on choice accuracy would increase over the course of the task, while the predictability of the 1D relative distance would stay the same.

The same cued goods were presented in both spaces. Following previous work on object and contextual novelty [39, 40], the experimental design distinguished context-related boundary changes (based on the manipulation of the boundaries in the space) from the associative structure related to the actual spaces(the contextual relevance of cued goods). Consequently for the cued goods, half of the cued goods inside the space maintained the same coordinates across abstract spaces (context-independent goods), while the other eight cued goods changed both their price and freshness variables depending on the space(context-dependent goods). Following the fMRI task, participants performed a surprise drag-and-rate task [41] on a laptop outside the scanner, where they were asked to place all the supermarket goods they previously saw on a blank map according to their 2D decision variable coordinates they most strongly associated with that particular good(Fig. 1D). We relied on this post-scan session to determine if participants retained a 2D representation of the decision spaces after the task.

Our analyses first focused on univariate fMRI signal differences related to the contextual relevance of a cued good versus the context type(type of shape) of its’ run during 2AFC trials. Ensuring that participants were representing 2D abstract knowledge in a similar way as previous studies[31], we tested whether the Euclidean distance between the cued goods to the landmark and closest boundary modulated fMRI representational similarity in the hippocampus and mPFC during the 2AFC. Testing whether hippocampus and mPFC signals were sensitive to geometrically changing abstract boundaries, we then used fMRI classifiers on boundary good encoding trials when decision variables weren’t visible to determine if there were distinct hippocampal-prefrontal representations for each boundary-defined abstract space. We could then relate each of these analyses back to choice and drag-and-rate performance. We expected hippocampal involvement in detecting contextual relevance and hypothesized that both mPFC and hippocampus would represent the Euclidean distance from the cued good to the nearest boundary good. Given the role of hippocampal cognitive maps in flexibly remapping their internal representation of the environment [9, 19–21, 24], we predicted that hippocampal fMRI signals would be sensitive to abstract boundary-defined contextual changes. To link any neural representations to behavioral performance, multivariate fMRI analyses were related to choice behavior, including the GLM choice accuracy analysis.

Here, we show participants build and maintain a 2D map-like representation of the two abstract spaces after the task. Asking whether changing boundaries in abstract spaces are represented in the hippocampus and mPFC, we observe that Euclidean distances between cued and boundary goods modulate fMRI pattern similarity in both the hippocampus and mPFC(Fig. 3), where only mPFC pattern similarity relates to individual performance improvements during the task. Moreover, we find that boundary-defined contextual identity can be accurately decoded from hippocampal fMRI signals with individual hippocampal classification accuracy relating to choice behavior.

## Results

### Behavioral results

The mean accuracy on the fMRI task was 81%(SD: 29%) and reaction time (RT) was 3.49s(SD: 1.79s), with participants responding significantly faster when they made correct decisions(t(28)=14.06, p*<* .001). To define the relative choice distance(RD), we computed the absolute difference between the relative distance of each good inside the space from the landmark and from the most proximal boundary for both decision variables (Relative choice distance = |ΔBoundary − ΔLandmark|). Participants’ accuracy significantly correlated with relative choice distance (*ρ*=0.23, p*<* .001; Fig. 2A), indicating that greater distances were less demanding. Similarly, RTs were faster when relative choice distances were greater(*ρ*=-0.14, p*<* .001). We then tested whether the type(contextual relevance) of cued goods or the boundary shape of the abstract space affected 2AFC task performance. We ran a 2x2 ANOVA to investigate the effects of boundary shape (square / distorted) and context dependency of the cued goods(context-dependent / context-independent) on choices. There was no significant interaction in between boundary shape and cued good type for choice accuracy(F(1,28)=0.3, p=.87) or RT(F(1,28)=2.99, p=.08). Furthermore, neither boundary shape (F(1,28)=0.81, p=.22 for accuracy and F(1,28)=0.53, p=.46 for RT), nor good type (F(1,28)=0.13, p=.72 for accuracy and F(1,28)=0.73, p=.39 for RT) influenced choice accuracy or RT (Fig. 2B). These results emphasize that participants behaved similarly during 2AFC trials independently of the contextual relevance of cued goods and the specific shape of the space where the trial occurred. To investigate whether there were changes in performance as the task progressed, we ran a 2x2 ANOVA for shape and repetition (whether it was the first or second time participants were making decisions in the shape). There was a significant effect of repeti-tion (F(1,28)=28.62, p *<* .001), but found no significant effect of shape(F(1,28)=2.61, p=.12) or interaction between shape and repetition(F(1,28)=0.32, p=.57; see Supplementary Figure 1A). This result highlights that participants exhibited learning-related improvements in task accuracy as the experiment progressed and performed equally well in both shape contexts.

Investigating if participants exhibited context-independent learning-related improvements in the task, we investigated the effect of 1D RD versus 2D Euclidean distance on choice accuracy. More specifically, we used a general linear model(GLM) to test whether trial-by-trial accuracy was significantly predicted by the relative 1D distance, the 2D Euclidean distance to the landmark good, or the 2D Euclidean distance to the closest boundary good. Crucially, this GLM also permitted us to measure whether the predictability of any of the three aforementioned variables on accuracy changed over the course experiment (Eq. 1). Consequently, this allowed us to distinguish general performance effects not linked to learning from ones related to learning. Only the Euclidean distance to the closest boundary both predicted accuracy throughout the task(t(28)=-4.03, p*<*.001) and significantly improved as the experiment progressed(t(28)=2.60, p=.015). The relative 1D distance also strongly related to choice accuracy(t(28)=4.65, p*<* .001), but unlike 2D distance its predictability didn’t improve as the experiment progressed(t(28)=0.961, p=.345; see Table 1 for group level statistics of all predictors). Notably, the lack of improved 1D RD prediction of accuracy as the experiment progressed couldn’t be explained by ceiling effects (Fig. 2B). These results are consistent with the hypothesis that participants build up map-like representations of multidimensional decision spaces that inform their choices, even when they aren’t needed.

**Figure 2:**
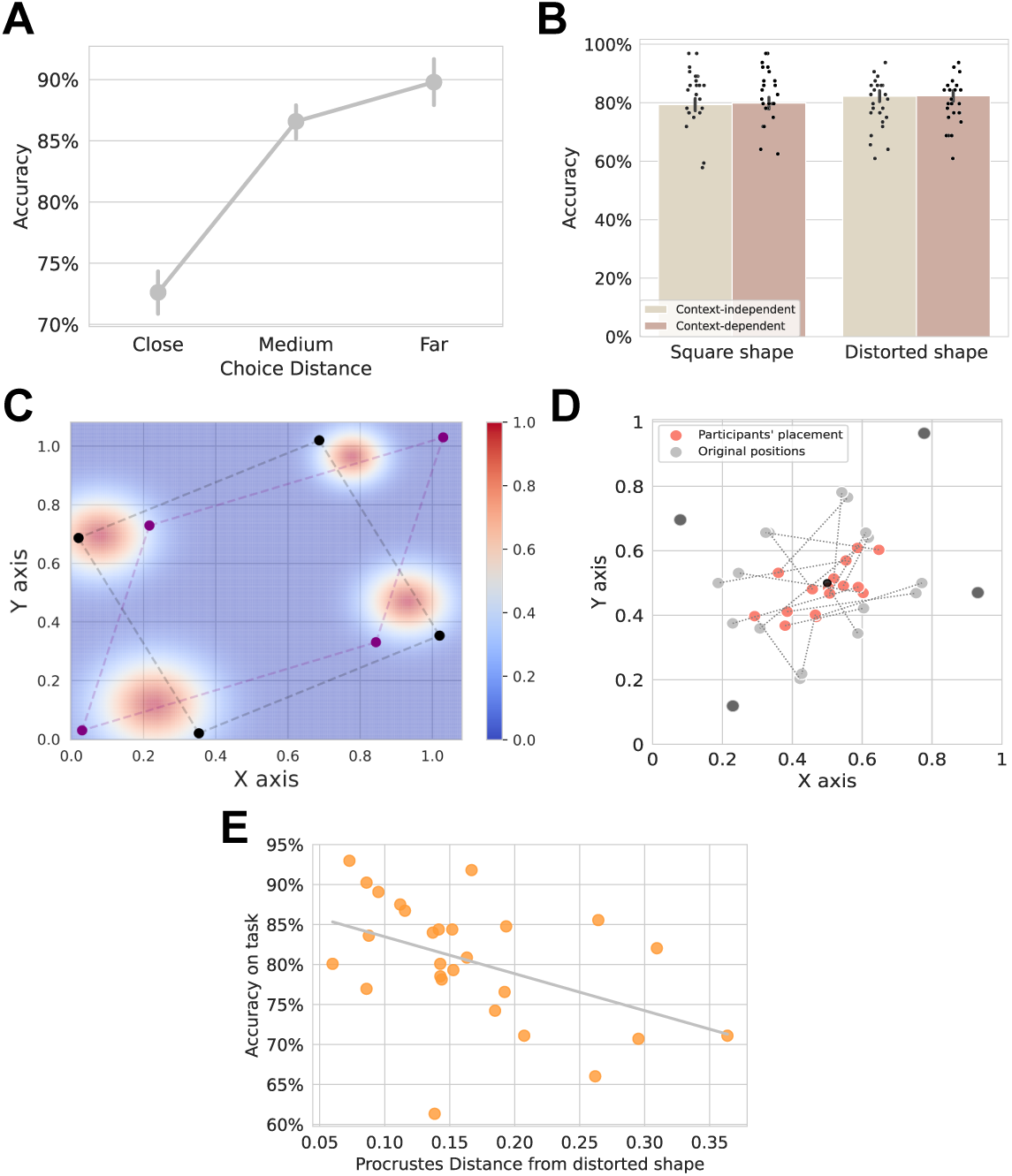
Behavioral results. **A**. Choice accuracy and relative choice distance. Participants’ choice accuracy significantly correlated with the relative choice distance of a decision trial (*ρ*=0.23, p *<* .001) Error bars represent SEM. **B**. Null effect of context shape and a good’s contextual relevance on choice accuracy. Each dot indicates a participant. Error bars represent SEM. **C**. Heatmap of mean boundary good placement. Warm colors represent standard deviation of participants’ placement of the borders, with the mean being the center. Square(black) and distorted(purple) shapes are overlaid for visualization purposes. Colormap reflects normalized distance units. Axes in D and E are normalized for visualization purposes. **D**. Post-fMRI cued good drag-and-rate placement across all participants. Pink dots represent cued goods according to participants’ reconstruction, gray dots represent the original position of the goods, and black dots represent participants’ mean boundary position across both shaped spaces. Dotted lines connect each cued good’s true and participants’ mean placement location. X and Y axes are normalized across price and freshness dimensions, where variables and axes were counterbalanced across participants. **E**. Correlation between choice accuracy and drag-and-rate task precision. Plot displays correlation between Procrustes distance of participants’ reconstruction from the original distorted shape with overall accuracy for all trials of the fMRI task (*ρ*=-0.42, p=.026).

**Table 1:** Group level T-Statistics Behavioral GLM.

| Predictor | T-statistic (28) | p value |
| --- | --- | --- |
| Choice Distance | 4.647 | < .001 |
| ED from Boundary | -4.027 | < .001 |
| ED from Landmark | -0.296 | 0.769 |
| Choice Distance × Run | 0.961 | 0.345 |
| ED from Boundary × Run | 2.596 | 0.015 |
| ED from Landmark × Run | 1.984 | 0.057 |

Further confirming that learning implicit 2D map-like representations of the different boundary-defined contexts informed participants’ choice behavior, we used a surprise post-fMRI drag-and-rate task to investigate if participants retained a detailed map-like representation of an abstract boundary-defined context. We computed the probability of placing the four boundaries in the specific sequences provided in the task(see Methods). Participants consistently organized boundary goods in the correct order(each boundary good shared a side with its neighboring boundary good) above chance (p*<* .001; Fig. 2C). We then examined whether the arrangement of all observed goods during the task accurately reflected the underlying 2D structure of the abstract spaces, thereby ensuring that all goods were positioned within the designated boundaries. We calculated the area occupied by the square and distorted shape in the space (45.1%) and computed the empirical probability of sixteen points falling within the boundaries of the space, which was p*<* .001. The overall percentage of cued goods placed within the boundaries across all participants was 68%, with a 20% variance (see Fig. 2D for mean placement).

We then tested whether participants reconstructed the boundary goods’ respective freshness and price coordinates in a proportional manner during the drag-and-rate task. Notably, participants’ reconstruction of the boundary goods was significantly wider along the freshness diagonal of the space(t(28)=6.7, p*<* .001), which is consistent with the freshness variable’s longer range(1-33 days compared to the price variable’s range of €1-€5.80). Confirming that the distortion of the freshness diagonal wasn’t due to decreased placement precision along that dimension, participants exhibited no significant difference in boundary placement errors along the price versus freshness axes (t(28)=-1.50, p=0.13).

Participants’ drag-and-rate behavior highlights they accurately placed the cued goods within the limits of the boundary goods. Crucially, post-task fMRI debriefing indicated that none of the participants were aware that the goods formed any kind of 2D space when performing the task. After confirming that participants placed the supermarket products within the boundaries above chance, we wanted to confirm that these effects weren’t due to spurious placement biases that didn’t reflect task learning. In particular, we wanted to rule out that participants clustered their placement of goods randomly around the center of the screen, or the center of the shape they constructed. To rule out this possibility, we compared placement distances from the center of the screen and shape with placement distances from the boundaries, where distance vectors were computed for each of the four boundaries from the sixteen cued goods for each participant. The minimum distance from a boundary was selected for each one of the cued goods and then compared to the distance from the center of the screen and the center of their created shape. Ruling out a centroid bias to the center of the drag-and-rate space, distances from the closest boundary were significantly smaller than distances from the center(t(28)=-2.36, p =.025). Furthermore, we observe that the minimum distance from boundaries is also smaller than distances from the centroid of the participants’ reconstructed shape(t(28)=-3.63, p=.002), thereby ruling out that participants randomly clustered around the middle of the shape they reconstructed. After we confirmed that participants retained a 2D representation of the decision spaces experienced during the task, we asked whether placement precision was related to individual differences in choice accuracy during the 2AFC session. We used Procrustes analysis to calculate the difference between the shape reconstructed by each participant (as defined by their boundary good placement) and the original coordinates of both the square and distorted boundaries. We then correlated the resulting Procrustes distance from each original space with fMRI task performance. The analysis showed that participants more faithfully reconstructed the original distorted shape in the drag-and-rate when they performed better on the decision task(*ρ*=-0.42, p=.026; Fig. 2E). However, a similar relationship wasn’t found for the square shape (*ρ*=-0.08, p=.70). Finally, we tested whether this effect could be driven by inconsistencies between the symmetry of the original square shape and participants’ reconstruction. We found that there was no significant correlation between the reconstruction of the mean shape coordinates(mean coordinates of square and distorted space boundary goods) and participants’ fMRI task performance(*ρ*=-0.36, p=.064). In addition, we correlated participants’ accuracy in a particular shape with Procrustes distance from that shape, observing no significant effect for either the square (*ρ*=-0.123, p=.541) or distorted (*ρ*=-0.271, p=.172) shape.

To further link the precision of participants’ 2D map-like drag-and-rate placement of the goods to the fMRI task, participants’ overall choice accuracy during the fMRI task was compared to participants’ ability to proportionally rescale the two dimensions of the boundary goods in the economic space. We calculated the difference between the diagonals on the x and y axis reconstructed by participants and related it to participants’ fMRI task performance, which yielded a significant correlation between participants proportionally rescaling the two drag-and-rate dimensions and individual fMRI task performance (*ρ*=-0.47, p=.017; see Supplementary Figure 1B). In other words, this indicates that participants who performed the memoryguided choice task better exhibited a smaller difference between the two diagonals in their drag-and-rate reconstruction(e.g., better rescaling across the two axes). In parallel, we analyzed participants’ placement of the landmark in relation to the centroid of their reconstructed shape. The distance of participants’ drag- and-rate landmark placement from the reconstructed centroid significantly correlated with individual fMRI task performance(*ρ*=-0.55, p=.003; see Supplementary Figure 1C), where participants who performed the memory-guided choice task placed the landmark closer to their reconstructed shape’s centroid.

#### fMRI analyses

##### Univariate

In a whole-brain fMRI analysis of 2AFC decision trials, we tested whether there were fMRI signal increases that related to the contextual relevance of cued goods and the shape of the abstract space. We ran a factorial analysis comparing differences between the four conditions(contextual relevance of cued goods versus the shape of the abstract space) in a GLM with parametric modulators for relative choice distance, accuracy, and RT. We observed a significant main effect of relative choice distance in the left posterior cingulate cortex, right fusiform gyrus, right precuneus, left lingual gyrus, right superior occipital gyrus, left superior occipital gyrus and left superior parietal lobule(see Supplementary Table 1 for more details). Examining the effect of accuracy on decision trials, there was a significant main effect in the left angular gyrus for making accurate decisions(see Supplementary Table 2 for more details). Additionally, we observed a main effect of faster RTs in right fusiform gyrus, left middle frontal gyrus, right precuneus, and right middle frontal gyrus(see Supplementary Table 3 for more details). Conversely, there was a significant main effect of longer RTs in decision trials in the right anterior cingulate cortex, left superior temporal gyrus, right insula, right supramarginal gyrus, right superior frontal gyrus, right inferior temporal gyrus, right middle frontal gyrus, left superior frontal gyrus, left cerebellum, right superior temporal gyrus and left insula(see Supplementary Table 4 for more details). However, there weren’t any significant effects of relative choice distance, accuracy, or RT in the hippocampus or mPFC. Contrary to our predictions, we observed no significant interaction between conditions in the hippocampus, nor anywhere else in the brain. The only significant accuracy-by-condition interaction we observed was in the right inferior occipital gyrus(see Supplementary Table 5 for more details), where the correlation with accuracy was higher in the distorted shape(t(25)=4.90, p=.004). There also wasn’t any accuracy-by-condition interaction in the hippocampus or mPFC ROIs. Furthermore, we didn’t observe any significant interaction in RT-by-condition or relative choice distance-by-condition in the hippocampus, mPFC, or anywhere else in the brain.

##### Multivariate RSA

We investigated whether the similarity of neural representations of the cued goods in the abstract spaces were modulated by their Euclidean distance to the landmark good and the most proximal boundary good. Euclidean distance was based on the pre-determined objective distance between the goods. Representational dissimilarity matrices(RDMs) featuring each cued good were created, resulting in a 16x16 matrix for each participant. The RDMs were included in a GLM where the predictors were behavioral RDMs based on experimental variables, including: a) 1D relative choice distance (RD), b) Euclidean distance of a cued good to the landmark, c) Euclidean distance of a cued good to the most proximal boundary good, d) contextual relevance of a cued good (context-dependent/context-independent; Fig. 3A). We observed a significant effect of Euclidean distance to the closest boundary in both bilateral hippocampus(t(25)=2.60, p=.015; mean explained variance: 35%. Fig. 3B, left) and mPFC masks(t(25)=2.39, p =.024; mean explained variance: 30%. Fig. 3B, right), suggesting that sharing a similar Euclidean distance to the boundary is a significant predictor of neural similarity across cued goods. We didn’t observe a significant relationship between representational similarity and Euclidean distance to the closest boundary anywhere else in the brain. Conversely, there was no significant effect of Euclidean distance of the cued goods to the landmark good in mPFC(t(25)=0.62, p=.54) or the hippocampus(t(25)=0.50, p=.62). Notably, we observed no significant difference between neural RDMs for Euclidean distances from the cued good to the landmark good and closest boundary good in the hippocampus (t(25)=0.88, p=.38) and mPFC (t(25)=0.80, p=.43). There were no significant effects for relative choice distance and representational similarity in the hippocampus(t(25)=1.70, p=.10), mPFC(t(25)=1.79, p=.09), nor any other brain region. Additionally, we didn’t observe significant effects for the contextual relevance of a cued good (i.e., whether a cued good’s 2D coordinates stayed the same between spaces) in the hippocampus(t(25)=-1.77, p=.09), mPFC(t(25)=-0.51, p=.61), or anywhere else in the brain.

**Figure 3:**
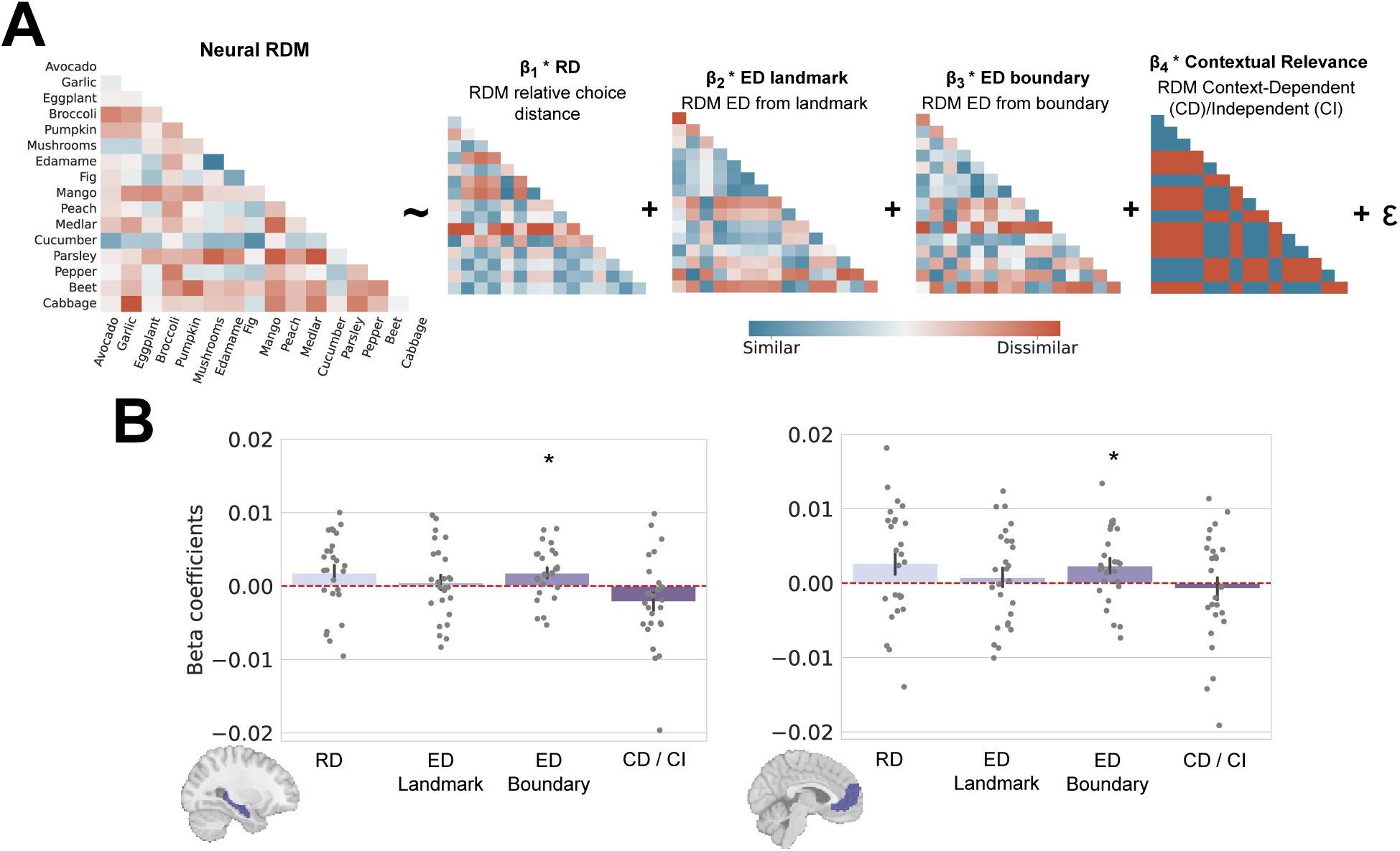
fMRI Representational Similarity Analysis(RSA). **A**. Representational dissimilarity matrices(RDMs) were computed by estimating neural dissimilarity from multivariate general linear models(GLM/neural RDM) and absolute difference of experimental variables(behavioral RDMs) of each pair of cued goods, resulting in 16x16 dissimilarity matrices. The behavioral RDMs were then entered as predictors in the GLM. Colder colors represent higher representational similarity. **B**. Beta coefficients of RSA GLM in hippocampus and mPFC. The bar plots show beta coefficients resulting from the GLM for each predictor; every participant is an individual Error bars represent SEM. Significant bilateral hippocampal effect of Euclidean distance from the cued good to the closest boundary good (t(25)=2.60, p=.015) and significant mPFC effect of Euclidean distance from the cued good to the closest boundary good(t(25)=2.39, p=.024). Asterisks showing significant effects: *= p*<* .05.

To rule out that semantic or visual similarity were driving the hippocampal and mPFC Euclidean distance from the nearest boundary effects, we ran control RSAs in the hippocampus and mPFC for both the visual and semantic similarity of cued goods. We didn’t observe significant visual or semantic similarity effects in the hippocampus (visual: T(25)=-1.44, p= .16; semantic: T(25)=-1.38, p=.18), or mPFC (visual: T(25)=.39, p=.70; semantic: T(25)=-1.55, p= .13). Furthermore, since cued goods that shared the same boundary in the 2D space were also presented with the same boundary during decision trials, we ran a control analysis to account for task-induced visual similarity. We computed trial-visual-similarity RDMs and correlated them with RDMs of hippocampus and mPFC, where neither region showed a significant effect(hippocampus: (t(25)=1.07, p=.29; mPFC: t(25)=0.79, p=.44; see Methods for further information).

To explore how neural representations of Euclidean distance relate to map-like learning, we then tested how hippocampal-mPFC Euclidean distance representations related to individuals’ Euclidean distance predictors of choice accuracy. We ran an inter-subject RSA (IS-RSA; see [42, 43]), for which we computed an intersubject RDM (nxn, n=26) comparing *β* coefficients of Euclidean distance to the closest boundary across participants for both the hippocampus and mPFC. We correlated these RDMs with a behavioral intersubject RDM, computed by comparing *β* coefficients of Euclidean distance to the nearest boundary from the behavioral GLM (see Supplementary Figure 2). Notably, individual mPFC RSA 2D distance effects related to individual 2D distance-related improvements in choice accuracy(*ρ*=0.180, p=.0011), which didn’t significantly relate to individual hippocampal RSA Euclidean distance effects(*ρ*=-0.087, p=.117). Consequently, these results link mPFC Euclidean distance representations to map-like learning of abstract knowledge.

##### Classifier analysis

Asking whether the hippocampus and mPFC flexibly represented abstract spaces consisting of the same goods with different boundary coordinates(price and freshness variables), we tested whether boundary-defined contextual identity(the shape of abstract spaces) was decodable from hippocampal and mPFC signals. We modeled fMRI data from all boundary good encoding trials when freshness and price variables weren’t visible to the participant(i.e., same visual stimuli in both abstract spaces). We conducted multivariate pattern analysis(MVPA) on encoding phase boundary good trials to test whether the patterns elicited during encoding trials in the square shape were distinguishable from trials in the distorted shape. We used a linear support vector machine(SVM), and applied a leave-one-subject-out cross-validation procedure(Fig. 4A) to test whether the classifier could decode the shape of the abstract space above chance by running one thousand permutations with shuffled labels between runs. In other words, training was conducted on N-1 participants to classify from hippocampal and mPFC ROI encoding trial data whether the remaining participant’s fMRI runs alternated between square and distorted contexts, or vice versa. The classifier yielded significant results in the hippocampus(p=.02, accuracy: 60%; Fig. 4B), but not in the mPFC(p=.29, accuracy: 53%). Furthermore, whole-brain MVPA searchlight analyses didn’t uncover any significant effects elsewhere in the brain. Testing whether classification of the abstract boundary shape was specific to boundary good trials in the hippocampus, we ran the abstract shape classifier on landmark good encoding trials and observed no significant effect in the hippocampus(p=.47, accuracy: 51%), suggesting that the hippocampal classification effect was specific to the representation of boundary goods. Additionally, there was no significant classification on landmark good trials in mPFC(p = .70, accuracy: 48%).

**Figure 4:**
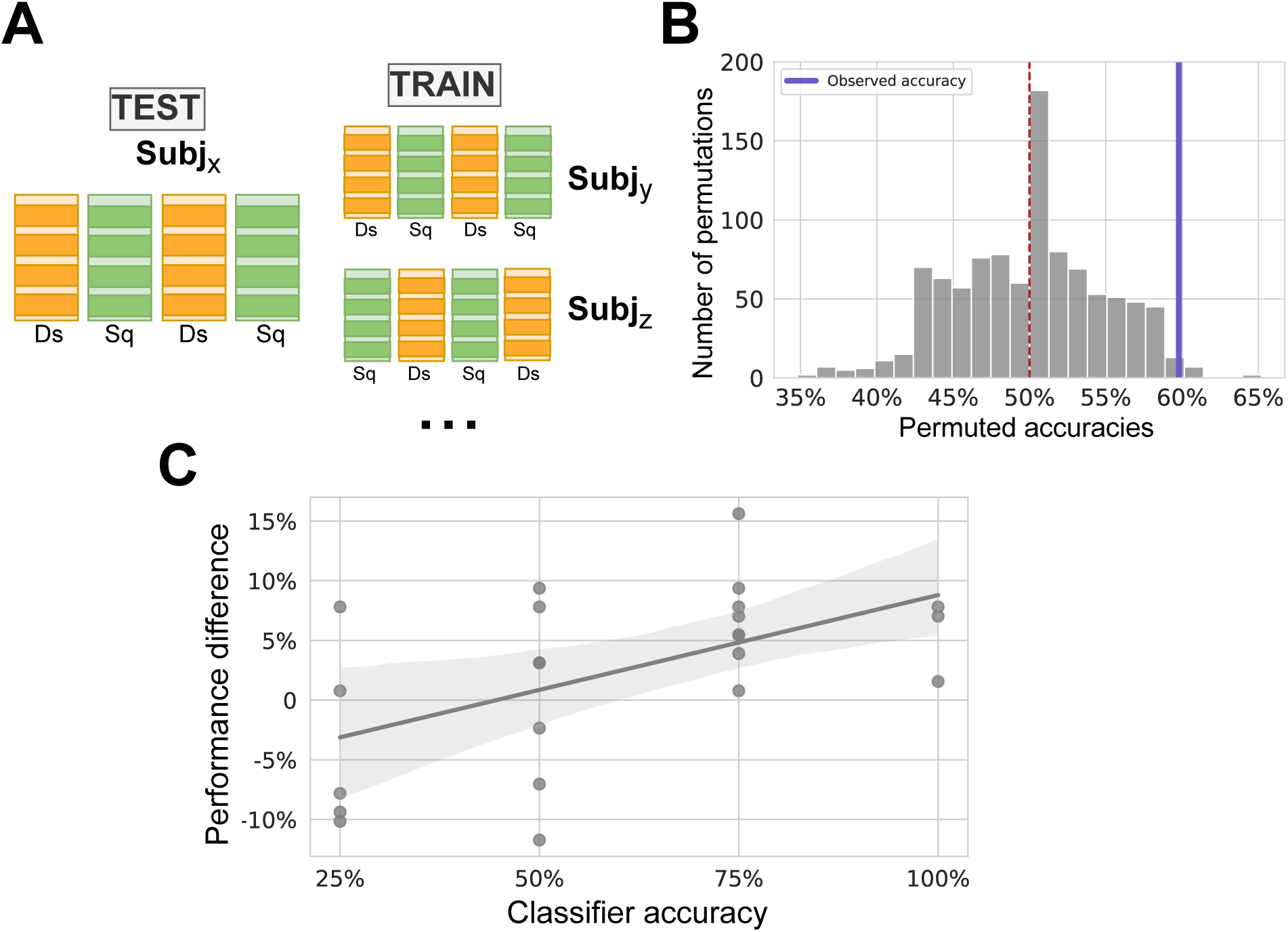
Hippocampal classifier analysis. **A**. Leave-one-subject-out cross-validation procedure. N validation folds were performed for N participants, where each validation comprised a classifier trained on N-1 participants and the Nth participant used as validation. In sum, training was conducted on N-1 participants to classify from their encoding trials(when price/freshness values weren’t visible) whether the remaining participant’s fMRI runs alternated between square and distorted contexts, or vice versa. **B**. Hippocampal classifier results based on one thousand permutations with chance level accuracy of 50%(indicated by red dashed line). Bold purple line indicates observed hippocampal classifier accuracy(p=.02, accuracy: 60%). **C**. Correlation between distorted versus square shape context task performance difference and hippocampal classifier accuracy(*ρ*=0.47, p=.022) for all boundary good encoding trials of the fMRI task. Positive differences indicate higher performance in distorted versus square shape.

The false discovery rate(FDR, e.g., the proportion of incorrectly classified observations for predicted class) of the hippocampal classifier was significantly higher for the square shape (Wilcoxon=29, p=.031). Given this difference, we ran a control classifier to check whether encoding trials in both shapes were classifiable above chance. Here, Gaussian noise was included as a third category and positive predictive values (PPV, e.g., the proportion of correctly classified observations per predicted class) on both the square and distorted shape were evaluated. Square and distorted shapes were accurately classified compared to noise(p=.00098 and p=.00014 respectively; see Methods and Supplementary Fig. 3 for further details).

Relating classifier accuracy to task behavior, we then asked whether hippocampal classifier accuracy related to participant choice accuracy. Classifier accuracy didn’t correlate with general task performance(*ρ*=0.22, p=.30). However, since the hippocampal classifier appeared to be biased towards the distorted shape, we then investigated whether the classifier bias towards the distorted shape corresponded with participants’ performance biases on the 2AFC task. Relating the difference in task performance in the distorted versus square shape to the hippocampal classifier bias, there was a significant correlation with classifier accuracy (*ρ*=0.47, p=.022; Fig. 4C) This result signifies that the classifier accuracy was highest in the hippocampus for participants that performed better in the distorted shape compared to the square shape.

Notably, this correlation wasn’t driven by performance effects related to an individual shape. There was no correlation between distorted context decision accuracy and hippocampal distorted context classifier performance(*ρ*=0.24, p=.261), nor between square context decision accuracy and hippocampal square context classifier accuracy(*ρ*=0.15, p=.502).

## Discussion

Using multivariate fMRI analyses and a novel memory-guided decision making paradigm, we asked whether changing boundaries in abstract spaces are flexibly represented in the hippocampus and mPFC. Participants built and maintained a 2D map-like representation of the two abstract spaces after the task(Fig. 2). We observed that Euclidean distances between cued and boundary goods modulated fMRI pattern similarity in both the hippocampus and mPFC(Fig. 3), where only mPFC pattern similarity related to individual performance over the course of the task. Moreover, we found that boundary-defined contextual identity can be accurately decoded from hippocampal fMRI signals, where classification accuracy is related to task performance(Fig. 4). In what follows, we relate our findings to the growing literature on hippocampal-prefrontal cognitive maps of abstract knowledge and discuss the cognitive implications of the hippocampus flexibly representing abstract boundaries.

We focused on hippocampus and mPFC following previous literature about the role of both these regions in forming cognitive maps and task representations [26, 27, 30–34, 36, 44]. Our results suggest that abstract boundary representations from decision spaces can be decoded from hippocampal fMRI signals. These results build on previous literature showing that neural ensembles in this region reorganize(remap) dynamically as a function of changes in the geometrical features of spatial environments, flexibly adapting cognitive maps in support of spatial memory [18–24]. Although we decode abstract spaces with different boundary representations in the hippocampus, it’s still unclear whether our results reflect hippocampal remapping computations. Furthermore, it’s important to note with our experimental design that we can only test how neural representational similarity and behavioral performance relates to the Euclidean distance between a cued good and its’ closest boundary good, not all four boundary goods. Consequently, it’s unclear how representing the Euclidean distance between the cued good and each boundary good influences choice behavior during the fMRI task. Some clues come from recent work that links category boundary-related memory distortions to hippocampal computations[45]. Future studies can build on this work, along with spatial paradigms investigating hippocampal remapping by gradually morphing abstract spaces or creating linear boundary changes [16, 17, 19, 20], to determine whether hippocampal spatial remapping computations are conserved in cognitive maps of abstract knowledge.

Previous studies on rodents suggest the mPFC could be a key candidate for building task representations, which allows for rapid generalization of spatial and abstract knowledge [39, 44].Similar findings in humans have highlighted the role of mPFC in supporting relational representations of task structure, even with different sensory events [37, 38, 44, 46]. Notably, we report mPFC involvement in representing Euclidean distances of task-relevant decision variables (Fig. 3B), while the coding of the boundaries doesn’t seem to be specific to the abstract space/context. This finding is consistent with ideas about mPFC involvement in organizing commonalities across experience [47]. A caveat to these findings is that we didn’t use an fMRI sequence that robustly measures signals in the most ventral portion of OFC, so these effects should be interpreted with caution.

Previous studies have shown how 2D cognitive map-like structure can be learned solely via inferred relationships [30, 31]. In contrast, we explicitly cued participants with both price and freshness decision variables during memory encoding, where they only made decisions on one dimension at a time. This is further re-inforced by our pre-fMRI training procedure, which involved participants encoding landmark and boundary good values immediately prior to making 12 forced-choices in five separate blocks. Consequently, it is unclear whether participants encoding the 2D coordinates for landmark and boundary goods facilitated cognitive and hippocampal-prefrontal map-like mnemonic representations. Still, our results show that participants retained a 2D representation of the task goods after decision making that reproduced the structure of the spaces, namely the boundary goods. Each cued good was paired with the most proximal boundary good, where participants placed some of the cued goods closer to other boundary goods while still keeping them within the limits of the space. This suggests learning of an implicit knowledge of task structure in the absence of explicitly learning cued goods’ coordinates relative to all four boundary goods [31, 35]. Moreover, participants reported no explicit awareness of the boundaries being positioned at the limits of the space and in a shape-like configuration when questioned after the fMRI task. Taken together, these results hint that even in the absence of extensive multi-day training, the hippocampus can learn and represent boundary-defined abstract contexts.

The question remains of why does the hippocampal classifier work significantly better on the distorted compared to the square shape context? We speculate that participants’ internal map-like contextual representations are partially distorted for both task contexts since participants don’t need to rescale the two differently ranged axis values to the same dimensions until after the fMRI task during the drag-and-rate. The idea of flexibly rescaling abstract map-like to current task demands meshes well with the idea that cognitive control processes can reshape the format of cognitive maps of abstract knowledge[43]. Developing new paradigms that use two different dimensions with the same range values without the need for rescaling, or more systematically change the range of dimensions, could be better suited to answer this question.

Our fMRI paradigm was designed to test for distinct neural processing for boundaries versus landmarks in an abstract 2D space, yet we didn’t observe significant changes in representational similarity with the Euclidean distance to the landmark in the mPFC, hippocampus, or other regions. Given the putative role of the striatum in landmark-based spatial navigation [21] and alignment in abstract knowledge spaces [48], it’s surprising that we didn’t observe any striatal involvement in representing the Euclidean distance to the landmark. One reason for this might be that unlike the aforementioned studies, participants make 1D proximity judgments instead of aligning the cued good coordinate to the landmark good. In our task, participants only need to know whether a cued good’s price or freshness is more proximal to the middle of the space or a boundary good, as opposed to whether the cued good’s value is higher or lower (or to the left or right) than the boundary good. Notably, proximity judgments are typically associated with boundary-oriented navigation [21]. Furthermore, the landmark was close to the center of both spaces, so it’s unclear how moving the landmark away from the center would change participant behavior. Future paradigms can move abstract landmark locations from the center and test alignment versus proximity judgments to more optimally compare landmark versus boundary learning processes in abstract knowledge spaces.

Segmenting the continuous flow of experience into distinct events plays a crucial role in episodic memory [49, 50]. Previous research has highlighted the hippocampus as a key region involved in event segmentation and the formation of event-specific [51–55] and context-specific representations [11, 12, 16, 20], which sometimes overlap [56]. Here, we relate hippocampal fMRI signals to more abstract boundary changes based on price and freshness decision variables of supermarket goods in abstract decision spaces. In contrast with spatial and event boundaries that share two primary dimensions, space and time[56], our results point to a more implicit use of boundaries distinct from explicit everyday experience. Given previous ideas about hippocampal cognitive maps also encoding episodic memory spaces [9, 24, 25], it’s unclear precisely how abstract boundary coding in the hippocampus might relate to hippocampal event boundary coding driven by spatiotemporal contextual shifts. However, there is evidence that expectations derived from spatial experience generalize to support temporal prediction[57]. Future research can investigate potential computational links between flexibly adjusting hippocampal representations of event and abstract spaces.

## Conclusion

We provide evidence that the human hippocampus flexibly represents abstract boundary-defined contexts, while mPFC representations more generally relate to learning 2D boundary-defined abstract contexts. Moreover, we find that participants learn these 2D boundary-defined abstract contexts during memory-guided decisions even when it isn’t necessary to effectively make a decision. Consequently, our data provide important clues on how the hippocampus and mPFC guide decision making across diverse spatial and abstract contexts.

## Methods

### Participants

34 participants (19 self-reported females and 15 self-reported males, mean age 23.4 years) recruited through an online questionnaire on Qualtrics (Provo, UT, USA, https://www.qualtrics.com) were remunerated and gave informed written consent to participate in the experiment. The study was approved by the local research ethics committee at Universitat Jaume I(ethics reference: CD/10/2022) and was conducted in accordance with the Declaration of Helsinki protocols. All participants were right-handed, had normal or corrected-to-normal vision, and had no history of neurological or psychiatric disorders. Of the 34 initial participants, 3 were excluded due to poor performance (task accuracy below 60%) and 2 due to technical issues with data collection. For the univariate and RSA fMRI analyses, we excluded an additional three participants due to excessive movement in the scanner and technical difficulties during image acquisition. An additional two participants were excluded from the MVPA analysis as they were missing a large portion of one run due to technical MRI issues, resulting in a sample of 24 participants for the MVPA analysis, 26 participants for univariate and RSA analyses, and 29 participants in the behavioral results.

### Task

We developed two abstract supermarket spaces to investigate memory-guided choices, where the two dimensions were represented by the price(in euros) and freshness(in days) of supermarket goods. Arbitrary price and freshness values were originally set to range from 1 to 33. To prevent the 2D shape configuration from being explicitly visualized by participants and present plausible prices, we then scaled down the price range to be between €1 and €5.80 by dividing the values by five. The four boundary goods were located at the extremes of the decision space in both square and distorted shaped contexts, changing their set of coordinates based on shape. The square shape had more extreme values on the price dimension, while the distorted shape had more extreme values on the freshness dimension. The landmark good was placed near the center of the decision space, maintaining the same coordinates in the square and distorted shaped contexts. All cued produce goods were located within the limits of the boundary goods. Each cued good was paired with a boundary good based on Euclidean proximity in the 2D space. Participants made similarity judgments on price or freshness for cued goods between the landmark and their most proximal boundary good. Additionally, eight of the sixteen cued goods inside the space maintained the same coordinates across abstract spaces(context-independent goods), while the other eight goods changed both of their positions(context-dependent goods). This experimental manipulation resulted in a 2x2 factorial design that tested the effect of boundary good shape(square/distorted) and cued good contextual relevance(context-independent/context-dependent).

The task was run using PsychoPy toolbox [58] on Python 3.9. Stimuli were supermarket goods overlaid on a gray background, and were viewed through MRI-compatible goggles. Pictures of all goods had the same resolution and were present in a 501x451 resolution frame. Participants were instructed that they’d make similarity judgments on the price and freshness of supermarket produce goods. The price and freshness of produce goods were random and not taken from real-world values in any local supermarket chain. Controlling for participants’ preconceived notions of price and freshness for different produce goods, we ensured there was no significant difference between how price/freshness values in the square and distorted spaces relate to real-life price and freshness values (price T(20)=-0.06, p=.95; freshness T(20)=0.12, p=.90; real-life produce prices taken from https://www.carrefour.es/supermercado and typical shelf life/freshness values taken from https://www.webstaurantstore.com/article/570/produce-storage-guide.html). Crucially, instructions didn’t mention the 2D configuration of abstract shapes or distinguish between the landmark and boundary goods. Prior to starting the fMRI task, participants underwent a training session on a laptop to ensure adequate task performance. The training consisted of an abbreviated version of the 2AFC task, featuring a novel abstract supermarket space with three boundary goods(a triangle shape), a landmark near the center, and six cued goods located within the limits of the boundaries. The training session lasted approximately fifteen minutes. We chose a training session of this duration to ensure that participants could effectively perform the task and efficiently make choices. We created a novel 2D triangle-shaped space during training to avoid priming participants towards one of the shapes in the fMRI experiment. After reading the instructions, participants were cued with both price and freshness variables for the boundary goods and the landmark good individually (“encoding phase”). They were instructed to passively view the goods and try to remember their price and freshness variables. Subsequently, decision trials on the six cued goods started(“decision phase”). Similar to the fMRI task, participants viewed a cued good on the top of the screen, while the landmark and the most proximal boundary were placed at the bottom. Participants had to decide whether the boundary and the landmark were more similar in price or freshness to the cued good. Participants were trained using a total of five blocks beginning with encoding and subsequently making twelve decisions per block. They received feedback on their accuracy at the end of each block. Importantly, since the training space only had six supermarket goods, the twelve decisions were repeated across the five blocks. To ensure understanding of the task, participants could see the relevant decision variable of the boundary and the landmark of each trial throughout the first block. If participants reached at least sixty percent accuracy on the last block, they proceeded to the fMRI task.

The fMRI task was structured in four separate runs, with participants performing one abstract shape per session with fMRI runs alternating between the square and distorted shape. The two shapes were the same throughout the task. The order of the sessions was counterbalanced across participants. Participants viewed the same task instructions as the training at the beginning of each fMRI run and were asked to press a button when they were ready to start. During the ensuing encoding phase, participants were cued with both price and freshness variables for each of the boundaries and the landmark individually. At each encoding trial, a picture of a boundary or landmark good was presented on the screen for three seconds, without any cue. Following a white fixation cross(jittered ITI, mean=2.5s), the good was presented again with its price and freshness variables for an additional three seconds(see Fig. 1A, “Encoding”). Participants were instructed to passively view the goods and try to remember their respective freshness and price variables. The presentation order of boundary and landmark goods was randomized. Participants were never cued on the shape they were performing during the run. After the encoding phase, participants started the decision making phase. At each decision trial, one of the sixteen cued goods inside the space was presented, with its relevant variable for the decision(price or freshness) being listed on the screen throughout the entire trial. At the bottom of the screen, the landmark and the most proximal boundary good were presented without any cued decision variables Fig. 1A, “Decision”). Participants were asked to choose which one between the boundary and landmark good was more similar to the given cued supermarket good. The decisions were made by choosing either one or the other by pressing the left or right button on the fMRI button box. Trials and positioning of correct choices were randomized across participants, where landmarks and boundaries left/right positions on the screen were counterbalanced. Trials timed out after nine seconds. Each of the cued goods inside the space had two types of decisions (“price” decisions and “freshness” decisions) for a total of thirty-two trial-unique choices per abstract space. During the fMRI task, participants never received feedback on their responses. To avoid excessive cognitive demands, participants were cued with the decision variables for the boundaries and landmark goods once they completed sixteen choices. Each run (shape) consisted of four blocks with interleaved encoding and decision periods. Participants made decisions for each cued good twice, resulting in sixty-four trials per run.

Following the fMRI task, participants performed a surprise drag-and-rate task from the Meadows web-based platform [41] on a laptop. They were presented with a blank map at the center of the screen with price and freshness index for x and y axes. (Fig. 1C). Axes were counterbalanced across participants, such that for some participants the x axis was indicated with “price” label and in others the x axis label was indicated with “freshness” label. Counterbalancing axes allowed us to control for axis anchoring to the origin during good placement. All supermarket goods from the 2AFC task were presented right next to the blank map, including the landmark good. Without time limitations, participants were asked to position each good according to the price and freshness coordinates(location) they most strongly associated with it.

### Choice behavior

All trials were included in the analysis of choice accuracy and RTs, where trials without a response were incorrect and assigned an maximum RT of 9s. We defined a relative choice distance for each trial, which was computed as the absolute difference between the distance of a given good from the landmark and from the boundary (Relative choice distance (RD) = |ΔBoundary − ΔLandmark|). We counterbalanced trials so that the mean relative choice distances for each condition–i.e. decision spaces, good types, and landmark/boundary proximity–were the same. Additionally, each of the cued goods inside the space had a landmark and boundary good as the correct answer depending on the trial (e.g, a good could be more similar to the landmark good in price trials and more similar to the boundary good in freshness trials), affording better coverage of the 2D abstract spaces. Additional variables were the Euclidean distance of the supermarket good from the landmark good and the boundary goods. All correlations were computed through Spearman’s rank correlation (*ρ*). To investigate whether choice accuracy could be predicted by 2D distances in the space, we ran a GLM for each participant including every trial:

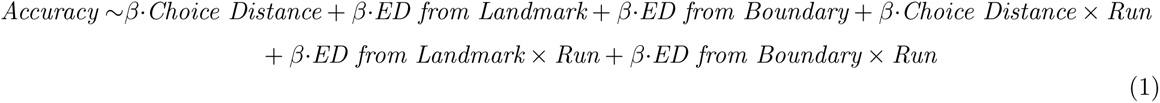

and then tested the beta coefficients for each predictor against zero for group-level statistics.

### Drag-and-rate

We quantified participants’ reconstruction performance in the post-scan drag-and-rate task. We computed the area of the screen occupied by the square and the distorted shape using the shoelace formula. Subsequently, we evaluated the empirical probability of 16 uniformly distributed points falling within these areas, which yielded a p *<* .001. To rule out a bias in clustering the goods’ placement around the center of the screen, we calculated the distance of every cued good from each of the four replaced boundaries and the center of the screen. We then selected the minimum distance from the boundary of each cued good. This resulted in two distance vectors for each participant, one reflecting distances from the boundary and one reflecting distances from the center of the space. We compared the two distance vectors with a paired t-test, measuring the magnitude of the difference between those distances for each participant. We tested the t-statistic from the paired t-tests against zero to calculate group level statistics. To investigate the different cued good placement from the boundaries and centroid of the reconstructed figure, we computed the centroid of the polygon resulting from individual boundary placement and then followed the same procedure.

Asking whether participants’ placement precision was related to individual differences in choice accuracy, we investigated participants’ ability to equally rescale the price and freshness ranges during the drag-and-rate task. Consequently, we computed the length of reconstructed diagonals by calculating the Euclidean distance between boundaries of each axis, which resulted in a “Price” and a “Freshness” diagonal. We ran a paired t-test to check for differences in the length of the two diagonals. Testing for individual differences in diagonal rescaling to equal dimensions, we correlated the difference in the length of diagonals for each participant with their fMRI task accuracy. We next investigated how participants’ placement precision of the landmark good. Since the landmark was originally close to the center of both square and distorted shapes, we asked how closely participants’ placed the landmark to the center of their reconstructed shape. Consequently, we computed the Euclidean distance between the centroid of each participants’ reconstruction and their replaced landmark, which was then correlated with their fMRI task performance.

### Procrustes analysis

Procrustes analysis was used to study the correlation between choice accuracy and drag-and-rate task precision, via the Procrustes library and Python 3.10 [59]. We quantified the similarity between the shapes in participants’ reconstruction by finding the best possible alignment/transformation between the matrices to maximize their similarity. This alignment involved scaling, rotating, and translating the data matrices to minimize the differences between corresponding points while preserving the overall structure.

### Permutation testing

First, testing data from the drag-and-rate task against chance distribution, we computed the probability of having five 2D points appearing in a specific sequence. The total number of possible sequences is 5!^2^, since we have five points in total and two coordinates. Therefore, for each permutation of the points in one dimension, there is a corresponding permutation in the other dimension. The probability distribution used is a discrete uniform distribution as each sequence of the five points has an equal probability of occurring, which means that the probability of each sequence is the same. The order in which the elements appeared in a sequence doesn’t affect their probabilities or outcomes. In this context, exchangeability suggests that any particular arrangement of the points is equally likely to occur as any other arrangement, provided that all points share similar characteristics and there is no intrinsic ordering among them. Specifically, the probability of any sequence is given by:

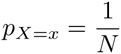

Where *X* is the random variable representing the sequence of five points, *x* is a specific sequence, and *N* is the total number of possible sequences.

### MRI Data acquisition

Whole-brain structural and functional MRI data were acquired using a 3T General Electric Signa Architect magnetic resonance imaging (MRI) scanner using a 32-channel head coil. Structural T1-weighted images were acquired using an MPRAGE sequence (TR = 2.5 s; flip angle = 8, and slice thickness = 1 mm). Four whole-brain functional runs were acquired using a T2*-weighted multi-band echo-planar imaging (EPI) sequences (voxel size= 2.5 mm isotropic, TR = 2 s; TEs = 20ms; flip angle *θ* = 90; resolution matrix = 80 × 80, multi-band acceleration factor = 2).

### Preprocessing

Data were processed within the fMRIPrep 23.0.2 framework [60]; which is based on Nipype 1.8.6 [61]. The T1-weighted image was corrected for intensity non-uniformity (INU) with N4BiasFieldCorrection, distributed with ANTs 2.3.3, and used as T1w-reference. The T1w-reference was then skull-stripped with a Nipype implementation of the antsBrainExtraction.sh workflow (from ANTs), using OASIS30ANTs as target template. Brain tissue segmentation of cerebrospinal fluid (CSF), white-matter (WM) and gray-matter (GM) was performed on the brain-extracted T1w using FAST (FSL 6.0.5.1). Brain surfaces were reconstructed using recon-all (FreeSurfer 7.3.2), and the brain mask estimated previously was refined with a custom variation of the method to reconcile ANTs-derived and FreeSurfer-derived segmentations of the cortical gray-matter of Mindboggle. Volume-based spatial normalization to one standard space (MNI152NLin2009cAsym) was performed through nonlinear registration with antsRegistration (ANTs 2.3.3), using brain-extracted versions of both T1w reference and the T1w template. For each of the functional scanning runs the following preprocessing within the fMRIPrep 23.0.2 framework was performed. First, a reference volume and its skull-stripped version were generated using a custom methodology of fMRIPrep. The estimated fieldmap was then aligned with rigid-registration to the target EPI (echo-planar imaging) reference run. The field coefficients were mapped on to the reference EPI using the transform. fMRI runs were slice-time corrected to 0.96s (0.5 of slice acquisition range 0s-1.92s) using 3dTshift from AFNI. The blood-oxygen-level-dependent (BOLD) fMRI signal reference was then co-registered to the T1w reference using bbregister (FreeSurfer) which implements boundary-based registration. Co-registration was configured with six degrees of freedom. Head-motion parameters with respect to the BOLD reference (transformation matrices, and six corresponding rotation and translation parameters) are estimated before any spatiotemporal filtering using mcflirt (FSL 6.0.5.1). Several confounding time-series were calculated based on the preprocessed BOLD signal: framewise displacement (FD), DVARS and three region-wise global signals. FD and DVARS are calculated for each functional run, both using their implementations in Nipype (following the definitions by [62]). The three global signals are extracted within the CSF, the WM, and the whole-brain masks. Additionally, a set of physiological regressors were extracted to allow for component-based noise correction (CompCor [63]). Principal components are estimated after high-pass filtering the preprocessed BOLD time-series (using a discrete cosine filter with 128s cut-off) for the two CompCor variants: temporal (tCompCor) and anatomical (aCompCor). The BOLD time-series were resampled into standard space, generating a preprocessed BOLD run in MNI152NLin2009cAsym space. The exclusion criteria for excessive movement was set with a FD greater than 0.20. The pre-processed BOLD time series data of each run were modeled with a general linear model (GLM) separately for univariate (using SPM 12) and multivariate analyses (using Nilearn 0.10.1 on Python 3.9, (https://nilearn.github.io/). In addition to task-relevant regressors, both GLMs contained twelve nuisance regressors, six of which pertaining to participants’ head motion (three rotation parameters and three translation parameters) and six pertaining to anatomical physiological noise (aCompCor). Moreover, BOLD fMRI data were smoothed with a 8 mm full-width half-maximum Gaussian kernel for univariate analyses, while no smoothing was performed for multivariate analyses.

### Univariate analysis

For each condition, we added parametric regressors based on relative choice distance, response times, and accuracy. This resulted in a *β* image of each condition of interest and its parametric modulators for each participant. For second-level analyses, we computed group level t-maps for each regressor and ran a 2x2 ANOVA(shape x cued good contextual relevance) based on our factorial design.

### Multivariate analysis

The preprocessed (unsmoothed) BOLD time series data of each run were modeled with a general linear model (GLM) on Nilearn using a boxcar function and the default canonical hemodynamic response function(HRF). For multivariate analyses, we ran two separate GLMs. In GLM1, we added regressors accounting for each of the sixteen supermarket goods inside the spaces during decision making. This resulted in a *β* estimate of activity for all trials of each good while participants were making decisions, on which we computed good-by-good RDMs. In GLM2, we accounted for timepoints in the encoding trials when price and freshness were not shown. A *β* estimate of activity was obtained for each boundary good, excluding visual differences between stimuli in the square and distorted shape.

### Representational similarity analysis

We ran representational similarity analysis (RSA) to investigate whether the behavioral variables in our memory-guided decision task relate to fMRI voxel patterns [64]. Representational dissimilarity matrices(RDMs) were compared across experimental conditions. We extracted voxels from hippocampus and mPFC for the *β* estimate of each good (GLM1), and computed RDMs of each cued good pair, resulting in a 16x16 neural RDM for each participant(Fig. 3A). RDMs of experimental variables were computed by calculating the absolute difference of that particular variable for each good pair, and entered those behavioral RDMs as predictors in a GLM where the dependent variable was the neural RDM. All RDMs were standardized before performing regression. The analysis resulted in a *β* coefficient of each predictor, for which we computed the mean of each participant and tested statistical significance against zero with a one sample t-test.

Linking individual behavioral GLM effects and fMRI RSA effects, we ran an inter-subject representational similarity analysis (IS-RSA; see [42, 43]). We computed an intersubject behavioral dissimilarity matrix (nxn matrix, where n=26) comparing *β* coefficients of Euclidean distance to the closest boundary from the behavioral GLM between participants. Next, we computed an intersubject neural dissimilarity matrix by comparing *β* coefficients of Euclidean distance to the closest boundary from the RSA GLM of hippocampus and medial prefrontal cortex ROIs between participants. We then correlated the two neural IS-RDMs to the behavioral IS-RDM.

To rule out potential confounds related to perceived similarity of the goods presented on screen, we ran a control RSA for both visual and semantic similarity of the cued goods. For visual features, we generated vector embeddings from the pictures of the goods used during the task (https://github.com/minimaxir/imgbeddings) and computed RDMs of their cosine similarity. For semantic similarity, we used a pre-trained word2vec model (https://github.com/piskvorky/gensim-data) to compute cosine similarity of word vector embeddings. We performed Spearman rank correlations between the resulting RDMs with neural RDMs of cued goods for every participant, and tested tatistical significance of the correlation coefficients on the group level with a one sample t-test. Furthermore, we ran a task-induced-visual-similarity control analysis on decision trials, which featured generated vector embeddings (https://github.com/minimaxir/imgbeddings) from snapshots of decision trials of each cued good and computed a visual-similarity RDM (vsRDM). First, we checked whether trials in which cued goods shared the same boundary showed greater visual similarity. We correlated the vsRDM with a model RDM that encoded cued goods that shared the same boundary, which yielded a significant result (*ρ*=0.61, p *<* .001). Next, we ran a partial correlation between the vsRDM and the neural RDM of both hippocampus and mPFC of each participant, while also controlling for the effect of spatial distances in the 2D space. Group-level statistics were conducted by comparing correlation coefficients against zero using a one-sample t-test.

In addition to ROI-based RSA, a searchlight-based RSA was conducted to investigate changes in neural pattern similarity related to 2D distances outside of our ROIs. We used *β* estimates of each good obtained from GLM1 and ran a four voxel radius searchlight analysis. We computed a neural RDM of the searchlight spheres based on each voxel of the brain [64]. Similarly to the ROI-based RSA, we then ran a GLM including the following predictor variables: a) relative choice distance; b) Euclidean distance of a good to the landmark; c) Euclidean distance of a good to the most proximal boundary good; d) contextual relevance of cued good (context-dependent/context-independent). The regression coefficients of the GLM for each searchlight sphere resulted in a whole-brain *β*-map of each predictor variable for each participant. In the second-level analysis, the regression coefficients were tested against zero using a one sample t-test. Analysis was performed using a Python open-source RSAtoolbox v.3.0 (https://github.com/rsagroup/rsatoolbox), and statsmodels toolbox v. 0.14.0 (https://github.com/statsmodels/statsmodels).

### Multivariate pattern analysis (MVPA)

We tested whether decision spaces with different shaped abstract boundaries were decodable from hippocampus and mPFC. A decoding analysis was conducted by using data from encoding trials of each shape and modeled data from the portion of the encoding phases when boundary good coordinates(price and freshness variables) weren’t visible to participants (GLM2). This resulted in a whole-brain *β*-map for each boundary in a run, averaged across encoding phases. Voxel patterns were extracted from the boundary *β*-maps in the hippocampus and mPFC ROIs. Since we were interested in decoding the activity from the different abstract spaces and not from the single boundary goods, mean signals were computed across the four boundary goods, obtaining mean patterns in ROIs for each run/shape. These patterns were included as features for the classifier. Asking whether we could decode which runs were square or distorted from encoding phase trial data in hippocampus and mPFC, two class labels of “square” and “distorted” were used for the four runs. We implemented a linear support vector machine(SVM) [65] with a principal component analysis(PCA) accounting for 90% of variance and applied a leave-one-subject-out cross-validation procedure [66]. Consequently, N validation folds were performed for N participants, where each validation comprised a classifier trained on N-1 participants and the Nth participant used as validation. This approach yielded a decoding accuracy value for each participant. For each run of each specific shape, the decoding accuracy values were constrained to discrete levels (0%, 25%, 50%, 75%, or 100%). The average validation classification accuracy across validation folds was used to characterize the expected decoding accuracy for the ROI. To check for differences in classification accuracy between square and distorted shapes, we computed the false discovery rate (FDR) of both shapes from the classifier confusion matrix. The FDR represents the proportion of incorrectly classified observations for predicted class, and is computed by calculating the ratio of false positives on all the positive calls (e.g, incorrectly predicted squares on all predicted squares). Testing whether the classifier could accurately decode above chance, we ran one thousand permutations with shuffled labels. Investigating whether the classification effect was exclusive to the boundary goods in the abstract spaces, we then ran the same classifier based on the landmark good, similarly extracting signals from our ROIs for when participants were looking at the landmark good (without price and freshness features) during the encoding phases of each shape.

To determine hippocampal classifier performance in distinguishing noise from the two distinct abstract space configurations, we implemented a linear SVM with a standard scalar preprocessing and ANOVA feature selection, selecting the 200 most responsive voxels in the hippocampus [65]. Data was augmented with a counterbalanced set of noise patterns by sampling and assigning two gaussian noise pattern vectors per participant for each cross-validation. We then computed the positive predictive value (PPV) for the square and distorted shape. The PPV represents the proportion of correctly classified observations per predicted class, and is defined by the proportion of true positives (TP) on all the positive calls; therefore, it gives information about the precision of the classification analysis in decoding each shape against the other and against noise. To check whether the decoding precision was above chance, we conducted five hundred label permutations by shuffling the square, distorted, and noise labels. These permutations were nested with five hundred validations corresponding to sampling a unique realization of gaussian noise pattern vectors.

To detect potential signals outside of our ROIs, we conducted searchlight-based MVPA, testing whether signals from square and distorted boundaries in other areas across the brain could be accurately decoded. The same procedure as the ROIs classifier was followed, using *β* estimates of each boundary obtained from GLM2. We applied a linear SVM kernel, PCA accounting for 90% of variance, and leave-one-subject-out cross-validation to searchlight spheres (radius=4 voxels) across each voxel of the brain. This resulted in whole-brain F-scores maps for which p-values were calculated. For the whole-brain searchlight MVPA classifier, Nilearn 0.10.1(https://nilearn.github.io/) was used.

### Statistical Analyses

Given our hypothesis, we focused our analyses on hippocampal and mPFC regions of interest(ROI) masks. We defined the bilateral hippocampal ROI using the Wake Forest University (WFU) Pickatlas [67, 68] integrated on SPM. The mPFC ROI was defined as the rostral medial wall of the prefrontal cortex, omitting ventral and frontopolar portions of mPFC, where robust signals weren’t observed in all participants. For univariate and RSA/MVPA searchlight analyses outside the two ROIs, we report effects surviving an uncorrected statistical threshold of p*<* .001 and correction for multiple comparisons(family-wise error correction, FWE) of p*<* .05 at the whole-brain level.

## Acknowledgements

We thank Doerte Kuhrt, Matthias Nau, and Ignacio Polti for helpful advice on multivariate fMRI analyses. We thank Michal Zareba for fMRI preprocessing advice. This research is supported by grants awarded to RK from the Valencian Community’s Program for the Support of Talented Researchers (CIDEGENT/2021/027), Universitat Jaume I Research Advancement Plan(UJI-B2022-45), and Spanish Science, Innovation, and University Ministry(PID2021-122338NA-I00).

## Contributions

Conceptualization, M.E. and R.K.; Methodology, M.E., L.A., A.G., and R.K.; Software, L.A. and A.G.; Formal Analysis, M.E. and L.A.; Investigation, M.E., L.A., and M.R.A.; Writing – Original Draft, R.K. and M.E.; Writing – Review & Editing, R.K.; Visualization, M.E. and L.A.; Supervision, R.K.; Funding Acquisition, R.K.

## Data and code availability

The data and code that support the findings of this study are available from the corresponding author upon reasonable request.

## Supplementary materials

**Supplementary Table 1:** Relative Choice Distance (RD)

| Region | x | y | z | Cluster Size | Cluster Level | T | Peak Level |
| --- | --- | --- | --- | --- | --- | --- | --- |
|  |  |  |  |  | p(FWE-corr) |  | p(FWE-corr) |
| L posterior cingulate | -2 | -38 | 39 | 120 | < .001 | 6.28 | .061 |
| R fusiform gyrus | 29 | -55 | -16 | 285 | < .001 | 6.2 | .074 |
| R precuneus | 6 | -58 | 24 | 533 | < .001 | 5.96 | .133 |
| L lingual gyrus | -12 | -78 | -9 | 404 | < .001 | 5.76 | .221 |
| R superior occipital gyrus | 31 | -78 | 22 | 159 | < .001 | 5.46 | .294 |
| L superior occipital gyrus | -24 | -90 | 19 | 109 | < .001 | 5.01 | .434 |
| L superior parietal lobule | -32 | -63 | 54 | 45 | .003 | 4.58 | .994 |

**Supplementary Table 2:**
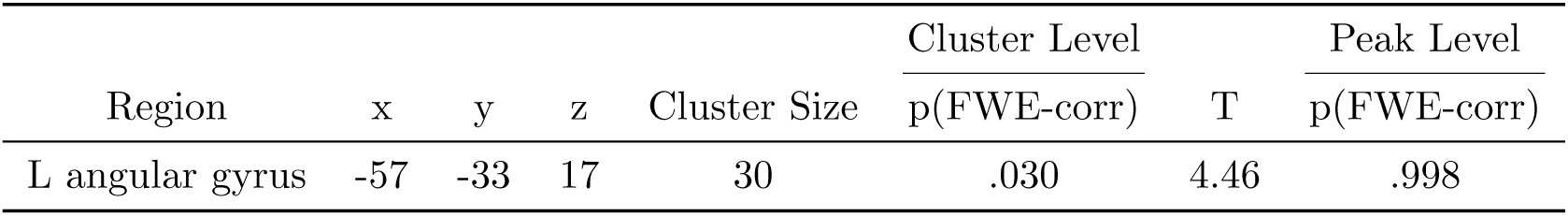
Accuracy.

**Supplementary Table 3:** Reaction Times -.

| Region | x | y | z | Cluster Size | Cluster Level | T | Peak Level |
| --- | --- | --- | --- | --- | --- | --- | --- |
|  |  |  |  |  | p(FWE-corr) |  | p(FWE-corr) |
| R fusiform gyrus | 41 | -70 | -11 | 5724 | < .001 | 9.68 | < .001 |
| L middle frontal gyrus | -27 | 0 | 57 | 42 | .018 | 4.68 | .952 |
| R precuneus | 1 | -55 | 42 | 67 | .001 | 4.65 | .961 |
| R middle frontal gyrus | 26 | 0 | 39 | 35 | .041 | 4.64 | .964 |

**Figure 4:**
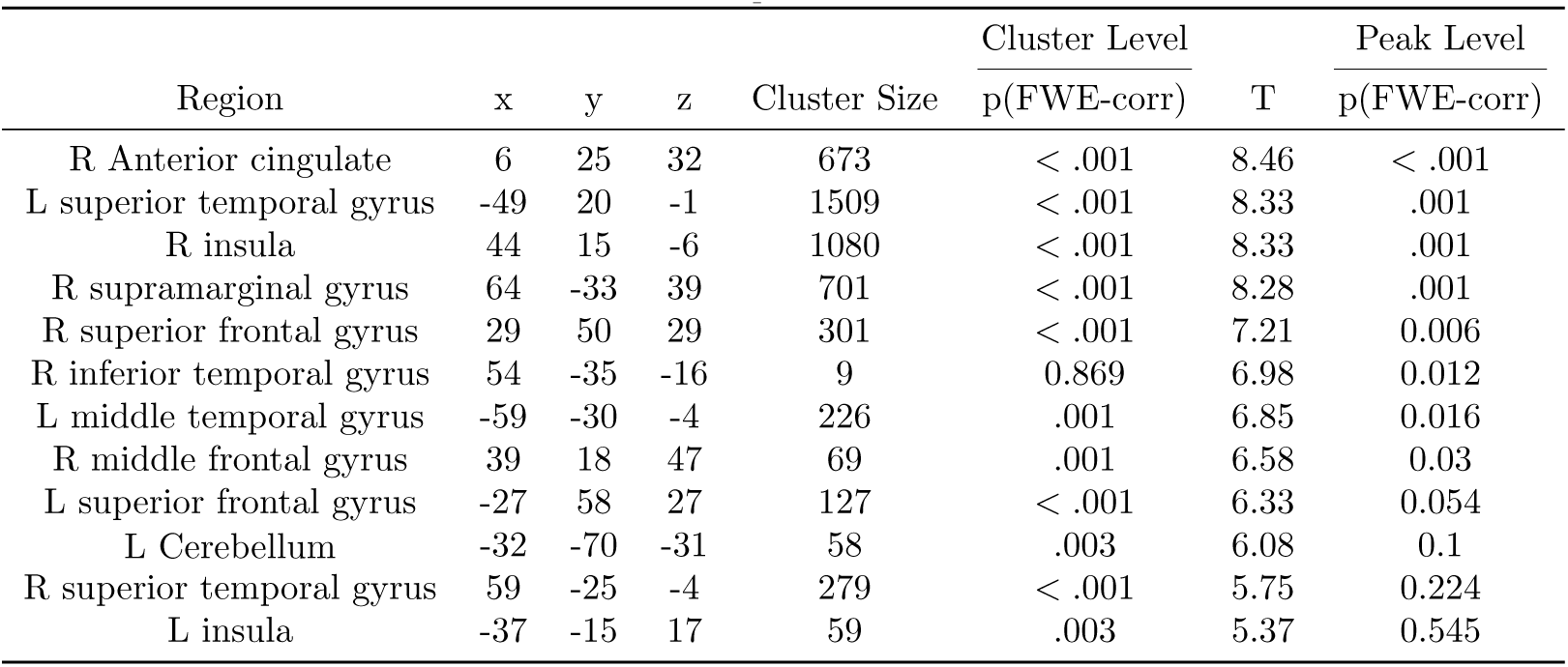
Reaction Times +.

**Supplementary Table 5:**
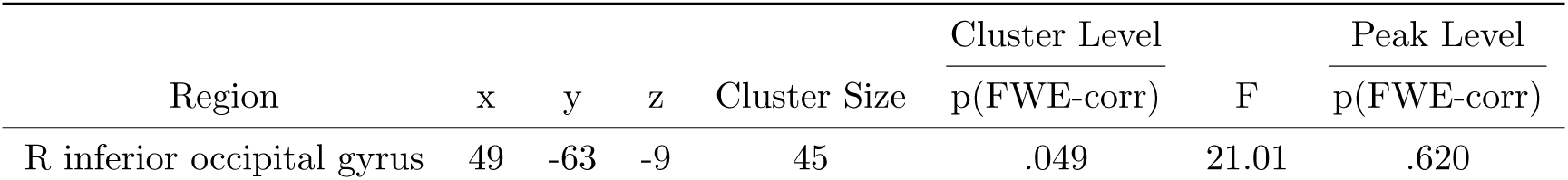
Accuracy By Shape.

| Region | x | y | z | Cluster Size | Cluster Level | F | Peak Level |
| --- | --- | --- | --- | --- | --- | --- | --- |
|  |  |  |  |  | p(FWE-corr) |  | p(FWE-corr) |
| R inferior occipital gyrus | 49 | -63 | -9 | 45 | .049 | 21.01 | .620 |

**Supplementary Figure 1:**
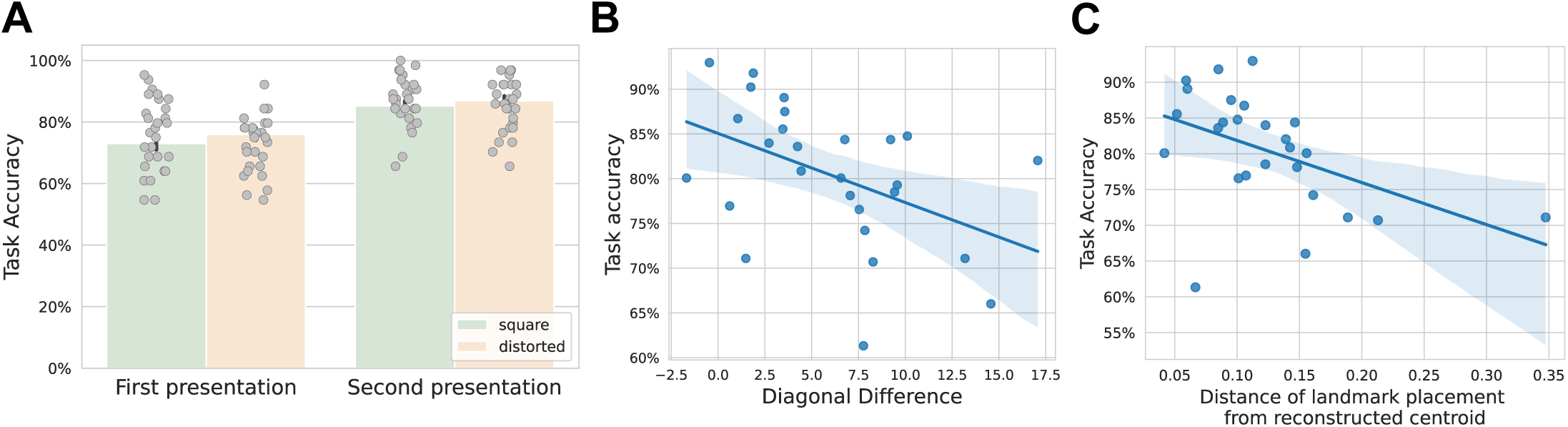
**A**. Significant difference in accuracy between first and second time participants see a shape, independently of shape identity. Each dot represents a participant. *** denotes p *<* .001. **B**. Significant correlation (*ρ*=-0.47, p=.017) between task accuracy and difference in diagonals of drag-and-drop reconstructed shape. C. Significant correlation (*ρ*=-0.55, p=.003) between task accuracy and distance of participants’ landmark placement from centroid of reconstructed shape during drag-and-drop.

**Supplementary Figure 2:**
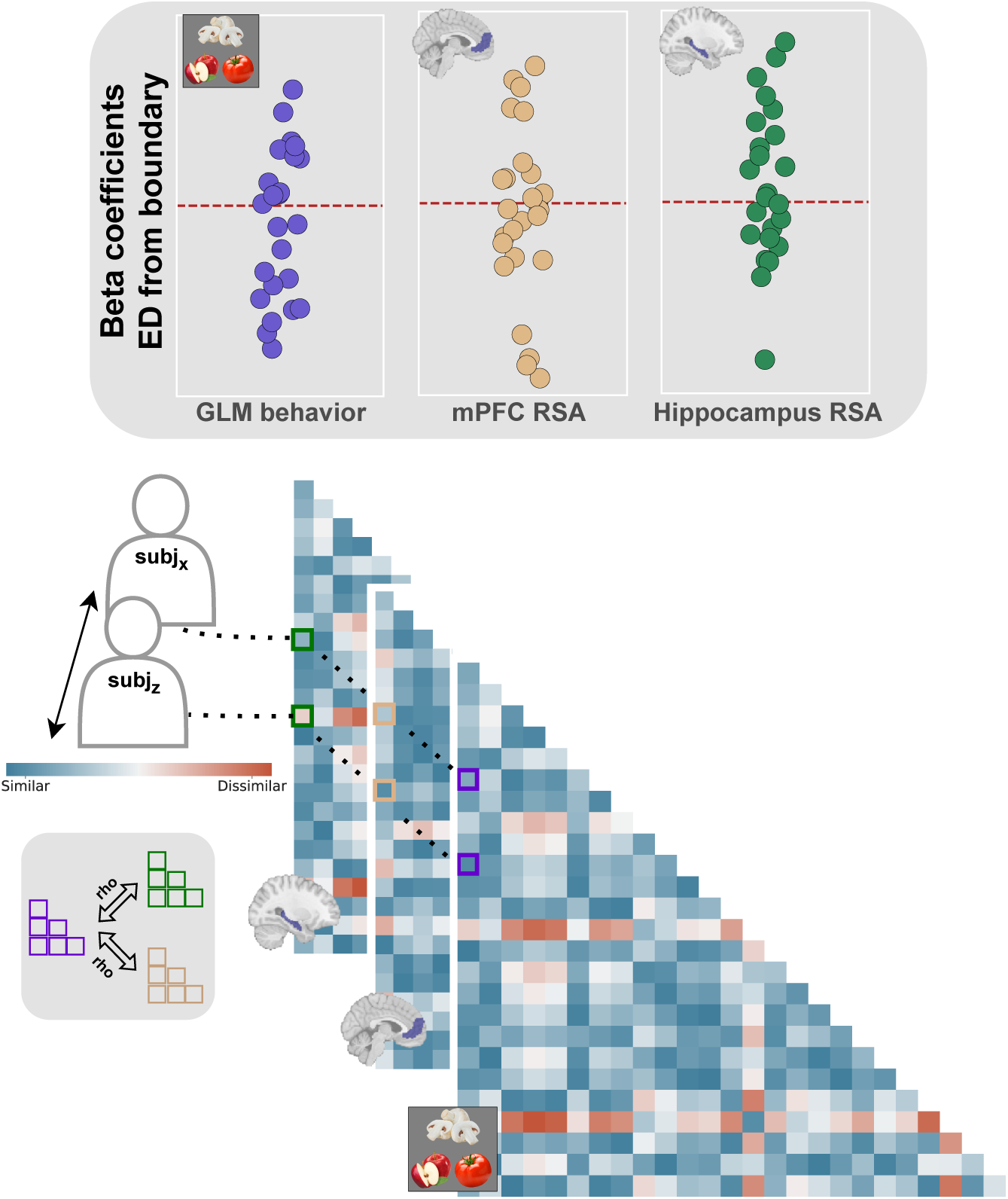
Graphical depiction of the Inter-Subject RSA methods. To investigate the relationship between 2D distance performance effects and neural representations, we conducted an inter-subject representational similarity analysis (IS-RSA). Specifically, we computed inter-subject representational dissimilarity matrices (isRDMs) based on the *β* coefficients from the behavioral GLM, which quantified the influence of Euclidean distance from the boundary on choice behavior. These isRDMs reflect the variability across participants in how ED affects behavior. In parallel, we computed separate isRDMs for the hippocampus and mPFC based on the *β* coefficients derived from the RSA GLM, which captured the neural response to the ED effect. These two neural isRDMs thus represent inter-subject variability in the neural encoding of the ED effect. Finally, to explore the relationship between behavioral and neural representations, we computed Spearman’s rank correlation(*ρ*) between the behavioral isRDM and neural isRDMs in both the hippocampus and mPFC.

**Supplementary Figure 3:**
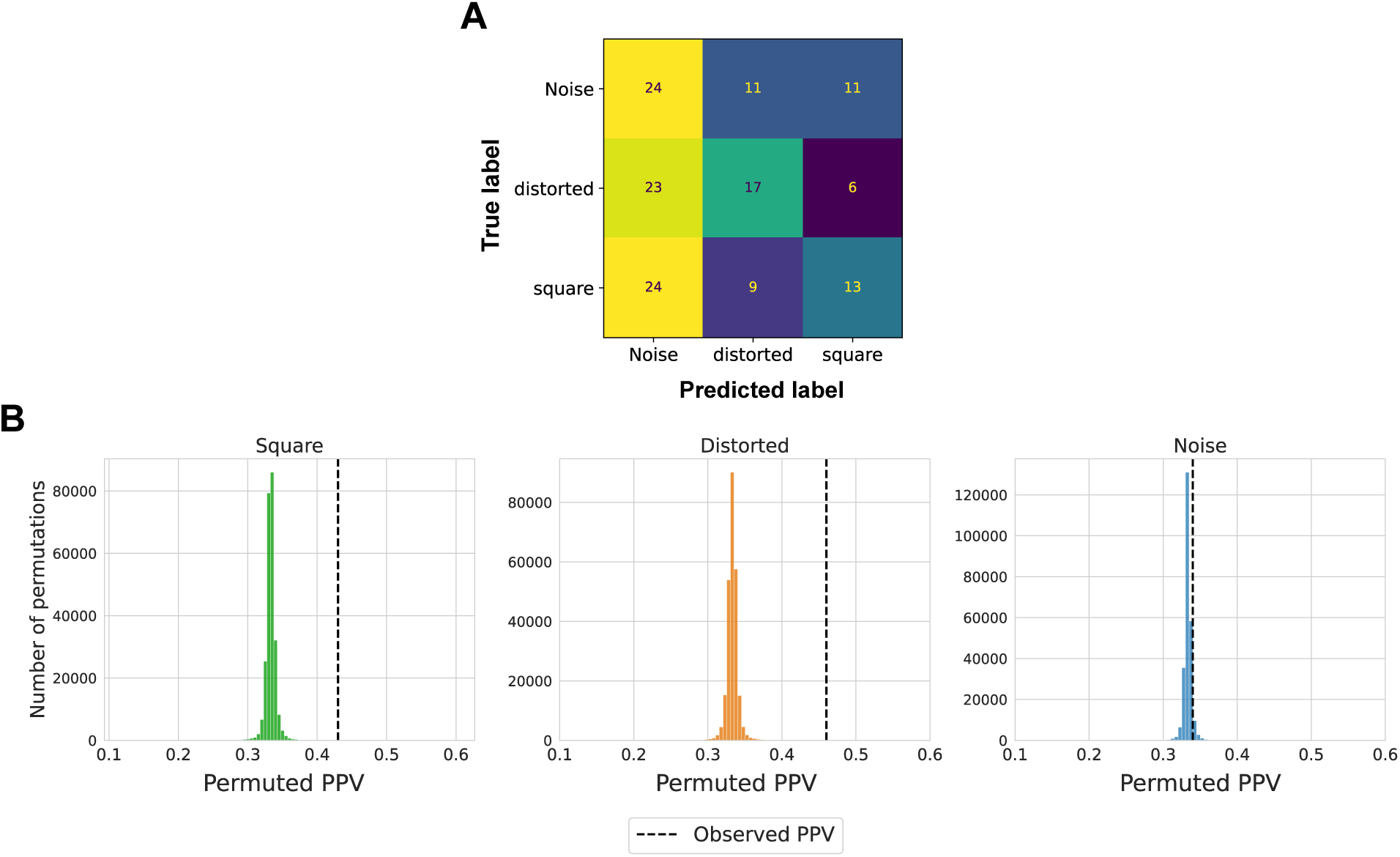
Control classifier analysis. **A**. Confusion matrix of the control classifier, with labels “Square”, “Distorted” and “Noise”. Noise pattern vectors were sampled from a gaussian distribution. **B**. Permuted PPVs of control classifier. We ran two hundred and fifty thousand permutations by nesting five hundred unique realizations of gaussian noise pattern vectors and five hundred permutations with shuffled labels. We then assessed whether classifier precision was above chance by comparing observed PPV (positive predictive value) with the null distribution. Plot shows PPVs obtained from permutations. Black dashed line indicates observed PPV (square = 0.43, distorted = 0.46, noise = 0.34).

